# Novel Small Molecular Compound AE-848 Showed Potent Inhibitory Activity Against Human Hepatoblastoma Cells

**DOI:** 10.1101/2022.09.12.507704

**Authors:** Juandong Wang, Qian Zhou, Likun Sun, Yaqi Xu, Yang Jiang, Xiaoli Liu, Shuo Li, Hui Li, Chengyun Zheng, Guitao Jie

## Abstract

**Purpose:** AE-848/33345007, is a novel small molecular compound that was independently synthesized by our laboratory. The aim of the current study was to investigate potential therapeutic effects of the small molecule compound AE-848 on human hepatoblastoma cells.

**Methods:** Human hepatoblastoma cell line HepG2 was cultured under different concentrations of AE-848. MTT assay was used to detect the inhibitory ability of AE-848 on the activity of this cell line, while flow cytometry was applied to quantify apoptosis. Western-blot technique was used to determine expression levels of signaling pathway-associated molecules. The antitumor *in vivo* activity of AE-848 against hepatoblastoma was evaluated using hepatoblastoma-bearing nude mice randomly divided into AE-848 (experimental) and saline (control) groups respectively. Changes in survival and tumor sizes were compared between these two groups.

**Results:** AE-848 significantly inhibited proliferation and induced apoptosis of HepG2 cells, and induced activation of PI3K/Akt/mTOR signaling pathways. More importantly, AE-848 administration significantly inhibited tumor growth and prolonged survival of hepatoblastoma-bearing mice.

**Conclusion:** AE-848 had a strong anti-hepatoblastoma activity both *in vitro* and *in vivo*, indicating the importance of developing AE-848 as a potential drug for hepatoblastoma treatment.

## Introduction

Hepatoblastoma (HB) is the most common embryogenic liver tumor in infants. It mostly occurs in infants under 3 years old, accounting for 80% of primary liver malignant tumors in children^[1]^**Error! Reference source not found.Error! Reference source not found.**. The disease is characterized by rapid development, rapid tumor growth, high degree of malignancy, and easy infiltration invasion and metastasis. At present, surgery combined with chemotherapy is widely used as multidisciplinary treatment^[2]^. Due to challenges with chemotherapeutic drugs, including postoperative residual, recurrence, intolerance, and drug resistance, one of the research hotspots has been to find safe chemotherapeutic drugs with low toxicity^**Error! Reference source not found**^.^[3]^. In this study, a novel small molecular compound AE-848^[4]^, is a novel small molecular compound that was independently synthesized by our laboratory. It was used to target the PI3K/Akt/mTOR signaling pathway. The effects of AE-848 on the proliferation and apoptosis of hepatoblastoma cells were determined, as well as the related molecular mechanisms of AE-848 to find new drugs and pathways for the treatment of hepatoblastoma. Administration of AE-848 significantly inhibited hepatoblastoma growth and prolonged survival of hepatoblastoma-bearing mice. These findings together provide experimental and theoretical basis for the therapeutic potential of AE-848 in preclinical studies conducted for the treatment of hepatoblastoma.

## Material and methods

### Reagents

AE-848: synthesized by our laboratory, chemical name: 5-bromo-2-hydroxy-m-benzaldehyde, molecular formula: C_24_H_21_BrN_8_O, molar mass: 517.39g/mol, chemical structure as shown in Figure **1**. AE-848 was dissolved in dimethyl sulfoxide (DMSO; Sigma, St. Louis, Mo, USA), with a concentration of 50mM and stored at 37 °C for use.

**Figure 1.**
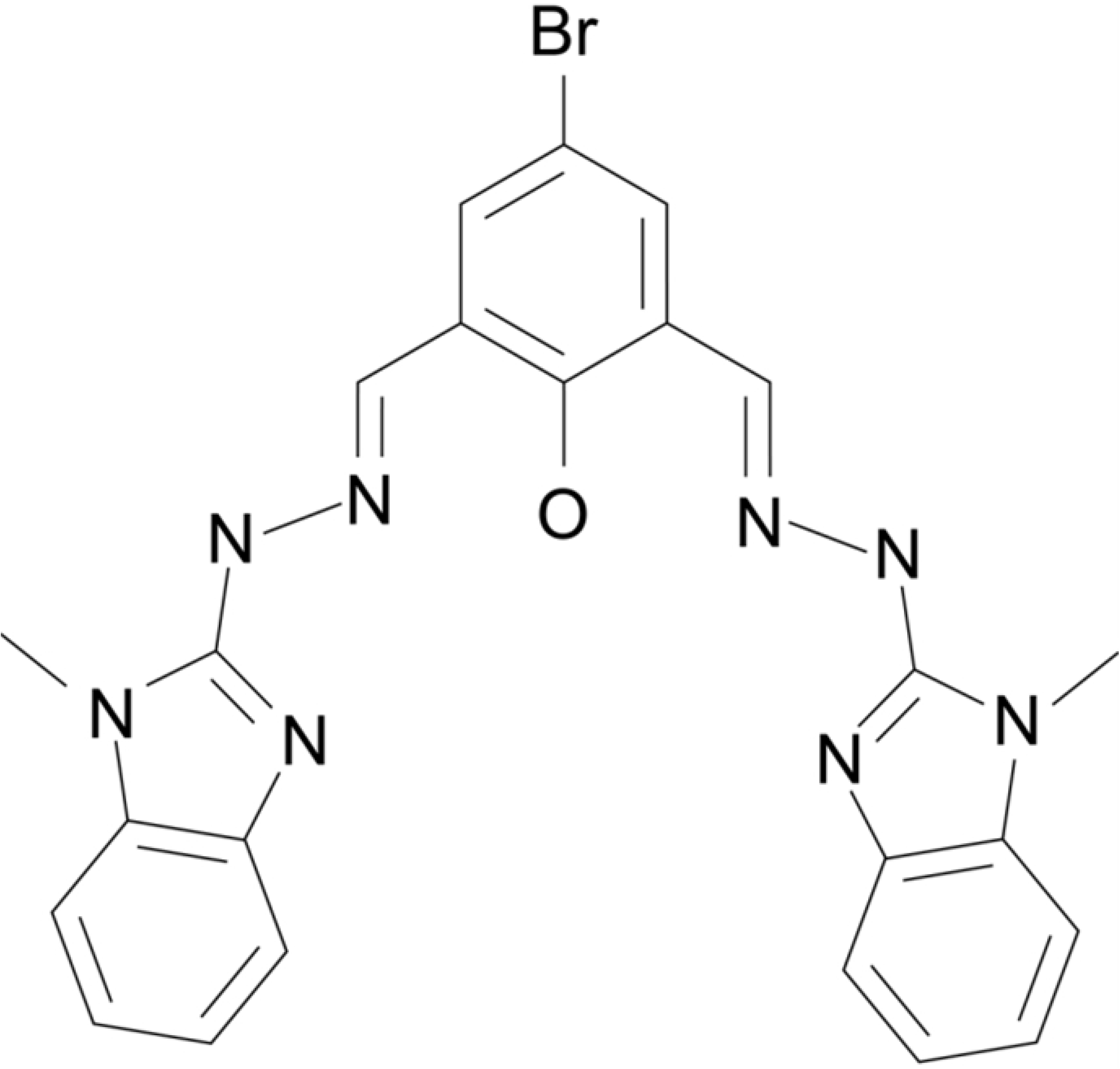
Chemical structure of small molecular compound AE-848

RPMI 1640 medium, fetal bovine serum, penicillin, streptomycin, and glutamine were obtained from Gibco (Waltham, MA, USA). Annexin V-FITC and propidium iodide (PI) test kits were purchased from KeyGEN BioTECH (Jiangsu, China) for detecting apoptosis. MTT was purchased from Solarbio (Beijing, China). BCA protein concentration assay kit and SDS-PAGE gel preparation kit were purchased from BOSTR (California, USA). Nuclear factor kappa B (NF/κB), phosphatidylinositol 3 kinase (PI3K), Akt, mammalian rapamycin target protein (mTOR), Bcl-2, Bax, GAPDH and β-actin were purchased from cell signaling technology (DANVERS, MA).

### Cell line

Human Hepatoblastoma HepG2 cell line was provided by the Central Laboratory of the second hospital of Shandong University. HepG2 cells were cultured in RPMI-1640 medium containing 10% fetal bovine serum (FBS), 100U/ml penicillin and 100mg/L streptomycin in a 5% CO2 incubator with a saturated humidity of 37 °C. In subculture, 0.25% trypsin was used for digestion.

### MTT cytotoxicity assay

Cell viability of HepG2 cells was determined by the MTT assay^[5]^. Briefly, the concentration of the HepG2 cells was adjusted to 1×10^4^ cells/well, and cells were treated with various AE-848 concentrations (2.5, 5, 10, and 20μmol/L) for 12h, 24h, 48h. At the indicated time points, 5mg/mL MTT was added to each well and incubated for 4h at 37°C, followed by decantation of the supernatant and addition of 10% DMSO to each well. The absorbance of each well was read on a microplate reader (Synergy Neo, BioTek, Winooski, VT, USA) at 570nm. To obtain significant experimental results, each procedure was repeated three times. Cell inhibition was calculated by the following formula: cell inhibition (%) = 100% – (average absorbance of treated group – average absorbance of blank) / (average absorbance of untreated group – average absorbance of blank) ×100

### Flow cytometric analysis of apoptotic cells

The apoptosis assay was performed as previously described^[6]^. Briefly, HepG2 cells were incubated in 12-well plates (1×10^5^ cells/well) with different concentrations of AE-848 (2.5, 5, 10, and 20μmol/L) for 12h, 24h, and 48h. Apoptosis was assessed using the Annexin V-FITC/PI kit (KeyGEN BioTECH, Jiangsu, China). The supernatant was collected, washed twice with PBS at 4°C, and digested with 0.25% trypsin without EDTA. Cultured cells were harvested by centrifugation and resuspended in the binding buffer, followed by Annexin V-FITC staining for 15min and PI staining for another 10min in the dark. Samples were analyzed with FACS Aria II (BD Biosciences, San Jose, CA, USA).

### Western blot analysis

HepG2 cells were cultured with various concentrations of AE-848 (2.5, 5, and 10μmol/L). Expressions of NF/κB, P-NF/κB, PI3K, P-PI3K, Akt, P-Akt, mTOR, P-mTOR, Bcl-2, Bax, GAPDH, and β-actin were analyzed by Western blot as described previously^[7]^. Briefly, drug-treated cells were lysed in RIPA buffer supplemented with a phosphatase inhibitor cocktail (Roche, Mannheim, Germany) and 1mM PMSF on ice for 20min. Protein concentrations were measured using the BCA Protein Assay kit according to the manufacturer’s instructions (Thermo Scientific Pierce). Whole protein (30μg/lane) from each sample was resolved on 10% SDS-PAGE, transferred to a PVDF membrane (Millipore, UK) and blotted overnight with various primary antibodies (NF/κB, rabbit polyclonal antibody, 1:1000, CST; PI3k, rabbit polyclonal antibody, 1:1000, CST; Akt, rabbit polyclonal antibody, 1:1000, CST; mTOR, rabbit polyclonal antibody, 1:1000, CST; P-mTOR, rabbit polyclonal antibody, 1:1000, CST; Bcl-2, mouse polyclonal antibody, 1:1000, CST; Bax, rabbit polyclonal antibody, 1:1000, CST; GAPDH, mouse polyclonal, 1:1000, CST; β-actin, rabbit polyclonal, 1:1000, Abcam), followed by a HRP-conjugated monoclonal secondary antibody (1:5000; Amersham Pharmacia Biotech, NJ) for 1h at room temperature. Membranes were exposed to the Immobilon™ Western Chemiluminescent HRP Substrate (Millipore, Billerica, MA, USA). Blots were detected on X-ray film using the ChemiDoc™ MP Imaging System (Bio-Rad).

### Xenograft experiments

All animal study procedures were approved by the Animal Ethics Research Committee of the Second Hospital of Shandong University. 20 SPF grade BALB/C nude mice (4~6 weeks of age, male, weighing 18 ~ 20g) were purchased from Jiangsu Jicui Yaokang Biotechnology Co., Ltd. (formerly Nanjing University model animal research institute). Under pathogen-free conditions, mice were fed in separate cages, and drank freely under suitable temperature and humidity, artificial darkness, and light alternate. When HepG2 cells grew to 80% ~ 90% confluence, they were digested with 0.25% pancreatin and centrifuged. HepG2 cells were made into a cell suspension, and the cell density was adjusted to 5×10^7^/ml using blood cell counting plate. Twenty nude mice were injected with 0.2ml cell suspension (1×10^7^) under the right lower extremity armpit. The cell suspension was fully mixed before injection. Approximately 7 ~ 10 days after inoculation, when the tumor grew to 3mm×3mm, patients were randomly divided into two groups. After inoculation, when the tumor diameter was more than 6mm, the tumor bearing nude mice with too large and too small tumor volumes were eliminated and the rest were randomly divided into two groups; normal saline and experimental groups, each group containing two sub-groups. One group was observed for the growth curve of nude mice, and the other group was observed for tumor growth. Observations included the general state of mice (body weight, eating, drinking water, *etc*.), measuring the body weight, the long diameter A and short diameter B of the tumor by calipers once every three days, calculating the tumor volume [V = 1/2 (AB^2^)], and starting administration when the tumor volume reached 100 ~ 150mm^3^. In the experimental group, AE-848 was injected intraperitoneally at a dose of 5mg/(kg·d) once a day for 2 weeks; in the control group, the same volume of saline was injected intraperitoneally once a day for 2 weeks. The mice were humanely euthanized by cervical dislocation when they reached the endpoint of the observation, which was defined as when the tumor size exceeded 2.0 cm in any direction or when a mouse was unable to creep for food and/or water. Changes in tumor volume and mice weight were monitored in the control and experimental groups every 3 days until the first mouse in the control group reached the endpoint.

### Statistical analysis

The differences between the groups were analyzed by the Student’s *t* test using SPSS 19.0. GraphPad Prism 5 (GraphPad Software Inc, San Diego, CA, USA) was used for statistical tests. *P* value < 0.05 was defined as statistical significance.

## Results

### AE-848 inhibited cell viability of HepG2 cells

AE-848 is a synthetic small molecular compound^[4]^ (Figure **1**). MTT assay was used to investigate the effect of AE-848 on viability of HepG2 cells. The inhibition rate of HepG2 viability gradually increased with elevated doses of AE-848. As shown in Fig. 2A, AE-848 was toxic to the HepG2 cell line (IC50-24h was 5.1 ± 1.8μM. To determine whether AE-848 had an effect on cancer cells in a time-dependent manner, the toxicity of one concentration of AE-848 (10μM) was tested on HepG2 cells for 12h, 24h, and 48h. As shown in **Error! Reference source not found.**, 61.5 ± 4.2%, 34.8 ± 1.8% and 20.4 ± 1.2% HepG2 cells survived after exposure to 10μM AE-848 for 12, 24, and 48 hours, respectively. Results showed that AE-848 inhibited the cell activity of HepG2 in a dose- and time-dependent manner.

**Figure 2.**
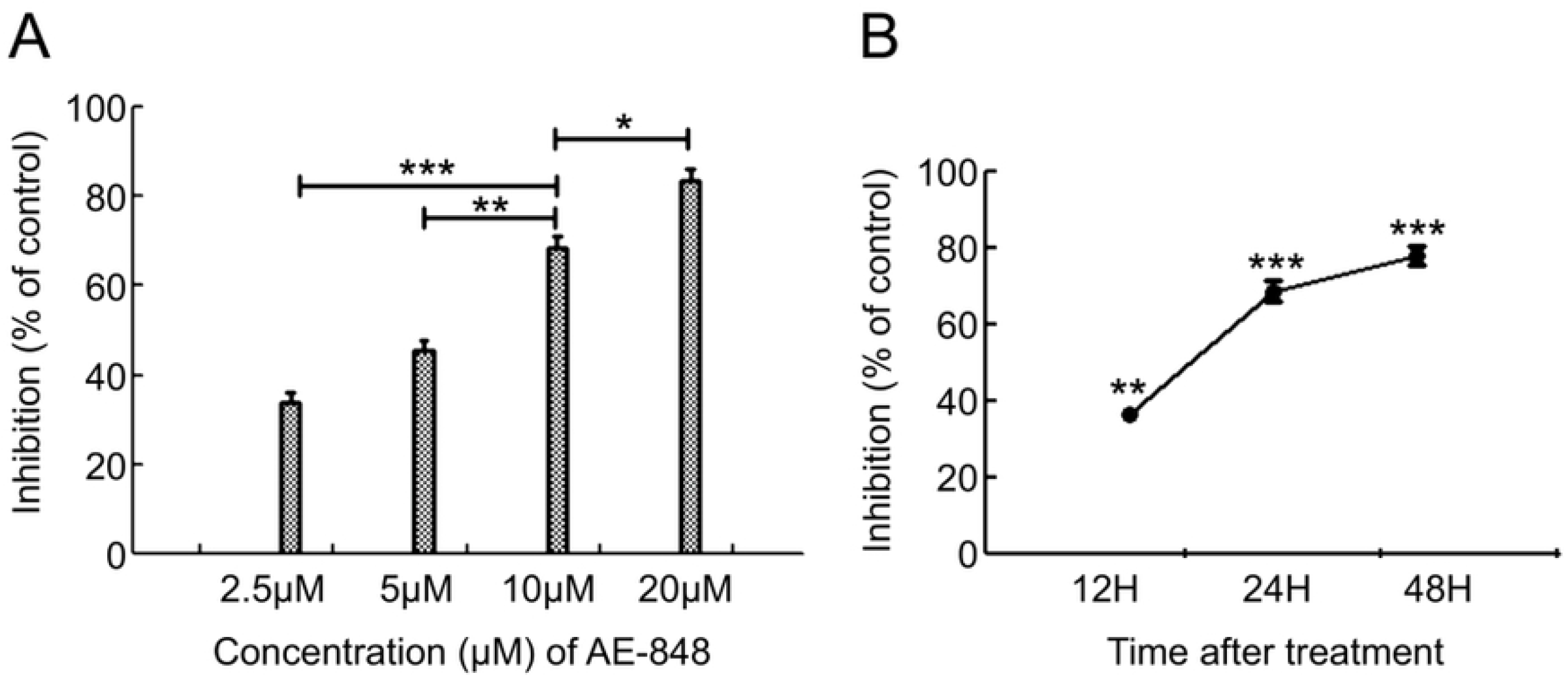
AE-848 inhibited human hepatoblastoma HepG2 cell viability in a dose- and time-dependent manner. (A) HepG2 cells were exposed to the concentrations of AE-848 (2.5, 5, 10, and 20μM) for 24h, after which the anti-viability effect was determined.

### AE-848 enhanced apoptosis of HepG2 cells

Apoptosis of HepG2 cells was detected by Annexin V-FITC and propidium iodide (PI) staining through flow cytometry. AV ^+^ PI^−^ and AV ^+^ PI ^+^ cells were counted as apoptotic cells. As shown in Figure **3**A-C, increased AE-848 concentration and the prolongation of exposure time increased the proportion of Annexin V positive cells. Compared with the control group (4.9 ± 0.86%), after treatment for 24h, the apoptosis rate of HepG2 cells was 7.5 ± 0.99% at 2.5μM, 15.9 ± 1.95% at 5μM, 52.5 ± 3.73% at 10μM, 80.2 ± 5.11% at 20μM, respectively (Figure **3**D). A similar pattern of change was observed in Figure **3**C-D, in which apoptosis increased significantly after incubation with 10μM AE-848 for 24 or 48 hours (52.5 ± 3.73% vs 4.9 ± 0.86%, *P* < 0.001; 93.6 ± 2.61% vs 6 ± 0.78%, *P* < 0.001). AE-848 induced apoptosis of HepG2 cells in a dose- and time-dependent manner, consistent with the findings in MTT analysis.

**Figure 3.**
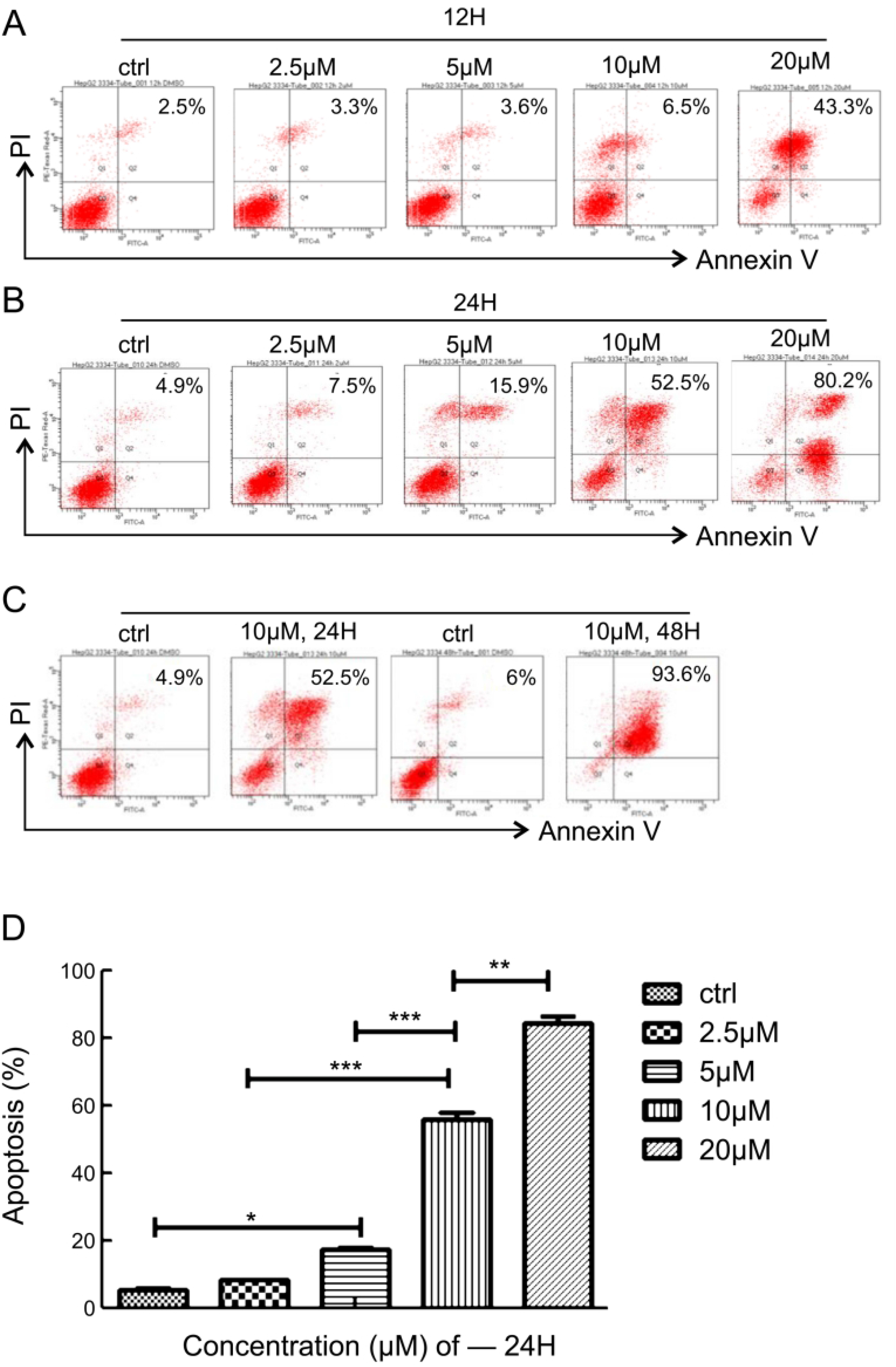
AE-848 significantly induced apoptosis in HepG2 cells. A, B, C) HepG2 cells were treated with a series of concentrations of AE-848 (2.5, 5, 10and 20μM) for 12h, 24h, 48h, after which the percentage of apoptotic cells was determined by flow cytom.

### AE-848 induced activation of apoptosis protein in HepG2 cells

Akt is a 56kD serine/threonine protein kinase, which is at the core of PI3-K signal transduction pathway. It is fully activated after phosphorylation under the action of PI3-K. Therefore, the phosphorylation level reflects the activation level of signal pathway. To prove that AE-848 inhibited PI3K/Akt/mTOR signaling pathway and induced apoptosis of HepG2 cells, Western blot was used to detect changes of Akt protein phosphorylation in HepG2 cells. Results showed that AE-848 inhibited Akt kinase phosphorylation in a dose-dependent manner, suggesting that AE-848 could inhibit the activity of PI3K/Akt/mTOR signaling pathway.

It was further investigated whether NF/κB and PI3K/Akt/mTOR signaling pathways are involved in AE-848-induced apoptosis of HepG2 cells. Bcl-2, Bax, NF-κB, PI3K, Akt and mTOR were detected. With increased AE-848 treatment concentration (2.5μM, 5μM, and 10μM) for 24 hours, the protein expression of NF/κB, p-NF/κB in HepG2 cells displayed no obvious differences (Figure **4**A); Bcl-2 levels decreased, and the protein expression of Bax increased noticeably (Figure **4**C), and phosphorylated PI3K, Akt and mTOR protein levels declined (Figure **4**A-B).

**Figure 4.**
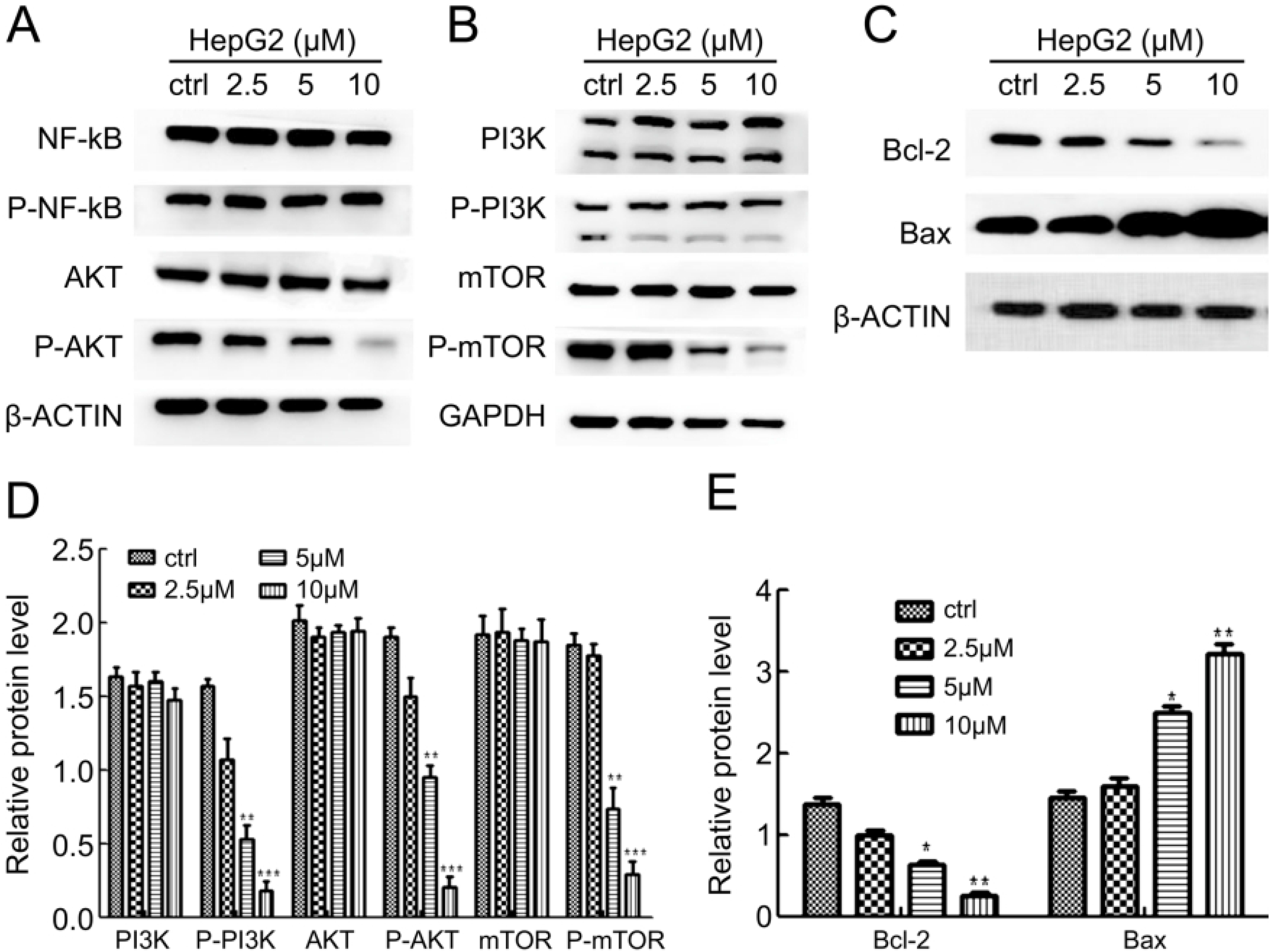
AE-848 inhibited the phosphorylation of Akt, PI3K, mTOR, and activated the apoptotic pathway of Bcl-2 and Bax. A) HepG2 cells were treated with various concentrations of AE-848 (2.5, 5 and 10μM) for 24h. Total proteins were extracted from the cultured cells and subjected to Western blot analysis using antibodies against NF/κB, Akt, P-Akt. GAPDH and β-actin were used as loading controls. B, C) Cells were treated in the same way described previously. Cell lysates were analyzed by western blotting to detect the expression levels of PI3K, P-PI3K, mTOR, P-mTOR, Bcl-2 and Bax. D, E) Shows the relative protein level of PI3K, P-PI3K, Akt, P-Akt, mTOR, P-mTOR, Bcl-2 and Bax calculated by the band density of western blots using Image J software of HepG2 cell line (\**P* < 0.05, \*\**P* < 0.01, \*\*\**P* < 0.001 by Student’s *t* test). Results were presented as Mean ± SEM from three independent experiments.

### AE-848 inhibits tumor growth and prolongs the overall survival of HB bearing nude mice

To investigate the therapeutic effect of AE-848 on human hepatoblastoma cell line HepG2 *in vivo*, AE-848 or saline containing DMSO and castor oil were injected intraperitoneally once a day for 14 days. As shown in Figure 5A-C, from the 7th day, AE-848 inhibited tumor growth. Regardless of tumor weight, volume, or size, AE-848 treatment group was significantly smaller than the control group (*P* < 0.01). Kaplan Meier curve was used to analyze the survival of the control and AE-848 group. The mean survival time of untreated control group and AE-848 treatment group was 17.0 days and 31.5 days, respectively (*P* < 0.01, Figure 5D). As shown in Figure 5D, AE-848 significantly prolonged the survival time of HepG2 tumor-bearing nude mice. AE-848 remarkably inhibited tumor growth and prolonged the survival time of HepG2 tumor-bearing mice.

**Figure 5.**
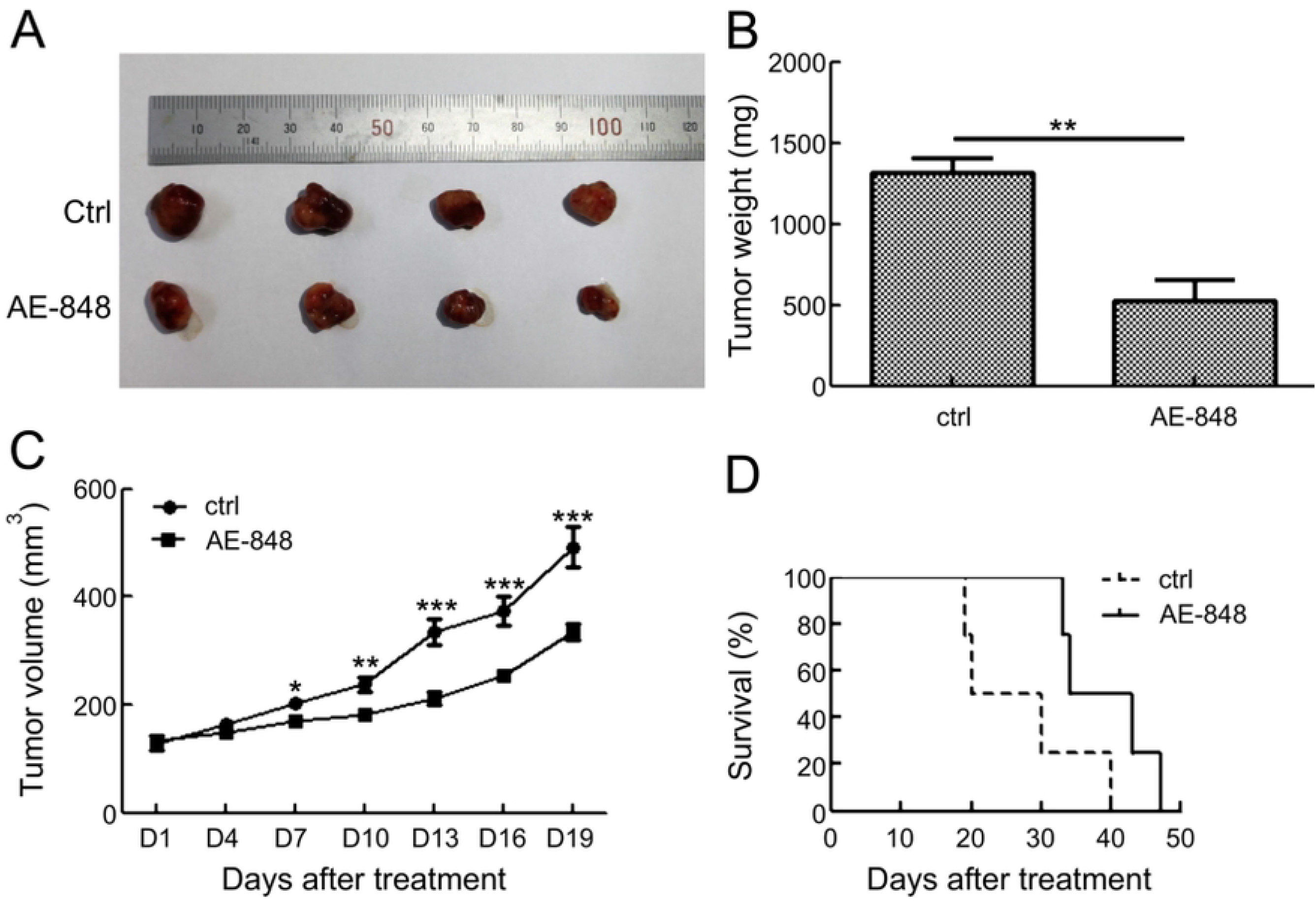
AE-848 significantly suppressed tumor weight, tumor growth and prolonged overall survival of hepatoblastoma bearing mice. AE-848 and control were administered to the hepatoblastoma-bearing BALB/C mice by intraperitoneal injection. A) Xenograft images were shown. B, C) Changes in tumor weight and tumor volumes among the control and treatment groups at the observation time points are indicated (\*\**P* < 0.01, \*\*\**P* < 0.001, by Student’s *t* test). The values are shown as mean ± SEM. Four mice in each group. D) AE-848 treatments significantly prolonged survival of the hepatoblastoma bearing mice as compared with the control (\*\**P* < 0.01, by Long-rank test, respectively). The values are shown as mean ± SEM. Four mice in each group.

## Discussion

AE-848/33345007, is a novel small molecular compound that was independently synthesized by our laboratory. Chemical name: 5-bromo-2-hydroxyisophthalaldehyde, molecular formula: C24H21BrN8O, molar mass: 517.39g/mol. In a previous study, this research group confirmed through *in vivo* and *in vitro* experiments, that AE-848 exerts obvious inhibitory effects on the proliferation of multiple myeloma cells^[4]^. To investigate whether the small molecule inhibits the proliferation of solid tumor cells, a human hepatoblastoma HepG2 cell line with higher incidence rate was selected.

As the most common primary liver malignancy in children, hepatoblastoma is a malignant tumor that originates in the embryo. It is usually diagnosed in children that are 1 – 2 years old. Its incidence rate in males is higher than in females. Over the past 60 years, the overall survival rate of hepatoblastoma has increased from 30% to 70%. This has been due to the implementation of standard chemotherapy, the improvement of surgical techniques, and the comprehensive application of preoperative transcatheter arterial chemoembolization^[8]^. Hepatoblastoma pathogenesis involves multiple factors and gene changes. At present, there is no widely recognized gene target or perfect molecular typing of hepatoblastoma. The specific etiology and pathogenesis of hepatoblastoma is unclear, and the diagnosis and treatment of hepatoblastoma need improvement. In addition to the occurrence of tumor genetic events, the occurrence, proliferation, invasion and metastasis of HB are also inseparable from various biological signals in the environment. The abnormal signal transduction pathway is also considered one of the pathogenetic pathways of hepatoblastoma. Recent studies have shown that, as a classic signal transduction pathway of promoting proliferation and inhibiting apoptosis, the PI3K/Akt/mTOR pathway plays an important role in promoting the growth and proliferation of tumor cells, inhibiting apoptosis, accelerating cell invasion and metastasis, promoting angiogenesis, and resisting cell apoptosis in chemotherapy and radiotherapy. Therefore, blocking the abnormal activation of PI3K/Akt/mTOR signal transduction pathway has become a new target of tumor therapy^[9–13]^.

PI3K/Akt/mTOR signaling is widely considered a central adjustment factor of cell proliferation, survival, and metabolism. A number of reports have shown PI3K/Akt/mTOR signaling to play a pivotal role in tumorigenesis, including HB^[14, 15]^. Some studies have demonstrated that the abnormal activation of PI3K/Akt/mTOR signaling pathway plays an important role in the occurrence and development of hepatoblastoma^[16, 17]^. Other studies have indicated that PI3Kα subunit mutation exists in hepatoblastoma, leading to increased kinase activity. *In vitro* inhibition of PI3K and subsequent reduction of phosphorylation of Akt and mTOR confirmed that the abnormal activation of PI3K/Akt/mTOR signaling pathway was key to the survival of hepatoblastoma cells^[16]^. For instance, tamoxifen down-regulating surviving expression in HB cell line HepG2 was mediated by PI3K/Akt/mTOR to induce apoptosis^[18]^. Li *et al*.^[19]^ claimed that downregulation of the PI3K/Akt/mTOR signaling could induce apoptosis in HB cancer HepG2 cells. Zhang *et al*.^[20]^also showed that PP7 could inhibit PI3K/AKT/mTOR signaling and induce autophagic cell death in HepG2. The activation of apoptosis is regulated by a variety of signaling pathways, among which PI3K/Akt/mTOR pathway is one of the most important^[21]^.

In this study, AE-848 concentrations were positively correlated with the expression levels of PI3K/Akt/mTOR signaling-associated molecules (Figure 4A/B/D). Overall, these results provide solid evidence of the involvement of AE-848 in HB progression through controlling the PI3K/Akt/mTOR signaling. After treatment with AE-848, the intracellular PI3K signaling pathway was effectively inhibited, cell proliferation was blocked and apoptosis induced, the proliferation of tumor cells was effectively inhibited, and the phosphorylation and activity of Akt were inhibited.

Bcl-2 and Bax are members of the Bcl-2 family that are closely related to the occurrence and development of apoptosis. They are the main regulatory factors of the mitochondrial apoptosis pathway, where they combine to form heterodimer complex, affect the permeability and membrane potential of mitochondrial membrane, jointly determine the changes of mitochondrial membrane potential, release relevant Pro apoptotic factors, initiate caspase cascade reaction, down regulate cell survival, and induce apoptosis^[22–25]^. In this experiment, AE-848 significantly increased the expression of Bax and decreased the expression of Bcl-2. In this study, with the increased AE-848 concentration and time of treatment, the inhibition of HepG2 cell activity also increased. Moreover, the activity of HepG2 cells was most significantly inhibited at the concentration of 10μM and 24h after treatment. These optimal conditions can serve as important parameters when designing subsequent clinical trials using AE-848 to treat hepatoblastoma patients. These results also demonstrated that the AE-848 induced apoptosis of HepG2 cells in a time- and dose-dependent manner. The molecular mechanism of AE-848 involved the inhibition of PI3K/Akt/mTOR signaling pathway and Bcl-2/Bax apoptotic pathway. Our results from animal studies also showed that AE-848 significantly inhibited tumor growth and prolonged overall survival *in vivo*, indicating a potent anti-hepatoblastoma activity of AE-848 *in vivo*. Further research entails investigating the role of AE-848 in other HB cell lines, and providing experimental and theoretical basis for preclinical studies of AE-848 in the treatment of hepatoblastoma.

## Conclusion

AE-848 induces apoptosis of hepatoblastoma cell line cells *in vitro* and *in vivo*, through its inhibitory effects on PI3K/Akt/mTOR signaling pathways, and activation of Bcl-2/Bax apoptotic pathway. Given its minimal toxicity to normal blood cells, which provides a therapeutic window, AE-848 is thus a promising candidate for the treatment of hepatoblastoma.

## Acknowledgments

This study was supported by the Key Research and Development Program of Shandong Province (No. 2019JZZY011115 and 2021CXGC011101), Weihai Zhengsheng Biotechnology Foundation and Rongxiang Regenerative Medicine Foundation of Shandong University (No. 2019SDRX-05). We would like to thank for Editage for English language editing.

## Author contributions

JW, CZ and GJ contributed to conceptualization, methodology, supervision, validation, and manuscript development. QZ and LS contributed to research and investigation including most cell culture experiments, data curation, and writing the manuscript. YX, YJ, XL, SL, QZ and LS contributed to the animal experiments. XL and HL contributed to research and investigation including cell culture, and formal analysis of results, writing, and editing the manuscript. All authors read and approved the final manuscript.

